# CRISPR-Analytics (CRISPR-A): a platform for precise analytics and simulations for gene editing

**DOI:** 10.1101/2022.09.02.506351

**Authors:** Marta Sanvicente-García, Albert García-Valiente, Socayna Jouide, Jessica Jabara-Wallace, Èric Bautista, Marc Escobosa, Avencia Sánchez-Mejías, Marc Güell

**Affiliations:** Department of Medicine and Life Sciences, Universitat Pompeu Fabra, Barcelona, Spain; Faculty of Mathematics and Computer Science, University of Barcelona, Spain; Integra Therapeutics S.L., Barcelona, Spain; Technical University of Denmark, DK-2800 Lyngby, Denmark

## Abstract

Gene editing characterization with currently available tools does not always give precise relative proportions among the different types of gene edits present in an edited bulk of cells. We have developed CRISPR-Analytics, CRISPR-A, which is a comprehensive and versatile genome editing web application tool and a nextflow pipeline to give support to gene editing experimental design and analysis. CRISPR-A provides a robust gene editing analysis pipeline composed of data analysis tools and simulation. It achieves higher accuracy than current tools and expands the functionality. The analysis includes mock-based noise correction, spike-in calibrated amplification bias reduction, and advanced interactive graphics. This expanded robustness makes this tool ideal for analyzing highly sensitive cases such as clinical samples or experiments with low editing efficiencies. It also provides an assessment of experimental design through the simulation of gene editing results. Therefore, CRISPR-A is ideal to support multiple kinds of experiments such as double-stranded DNA break-based engineering, base editing (BE), primer editing (PE), and homology-directed repair (HDR), without the need of specifying the used experimental approach.

## Introduction

CRISPR-based gene editing has become a fundamental toolbox to cover a large variety of research and applied needs. It facilitates the editing of endogenous genomic loci and systematic interrogation of genetic elements and causal genetic variations^1–3^. Nowadays, it is even on the verge of becoming a therapeutic reality *in vivo*^4^. Despite tremendous advances, DNA editing and writing still involve imperfect protocols which need to be optimized and evaluated. This makes it essential to have tools that enable accurate characterization of gene editing outcomes.

Gene editing outcomes often involve complex data sets with a diverse set of genotypes. This is especially accentuated for double-stranded DNA-based gene editing such as those based on non-homologous end-joining (NHEJ) or homology directed repair (HDR). These experiments often generate complex gene editing signatures involving insertions, substitutions, and deletions.

Accurate quantification of this distribution of genotypes may have important implications including knockout integrity or splicing modulation. On one hand, experimental conditions, gene editing reagents, and cell types influence gene editing outcomes. Analytical tools, and simulations are needed to help in the experimental design. DNA repair outcomes have incomplete predictability^5^, and deep exploration of DNA repair pathways shows that double strand break (DSB) repair gene mutations with similar sequences can come from different repair mechanisms^6^. Unraveling this complexity can help in the development of editing tools with higher efficiency and specificity, as well as in the implementation of better prediction models. In addition, when certain genotypes have to be enriched, prediction is even more relevant in design. One example is HDR improvement by targeting indel byproducts, where the prediction of the outcomes of the first target edition, allows the design of a second one to give a second chance to HDR^7^. On the other hand, a precise gene editing assessment is essential for validation purposes.

Initial methods for editing assessment were based on T7 endonuclease 1 (T7E1) mismatch detection assay and Surveyor Mismatch nucleases enzyme assays, which are based on the detection of DNA heteroduplexes after PCR amplification, denaturalization, and reannealing. When editing takes place, the different reanneled products generate heteroduplexes, which are cleaved by T7E1^8^. Tracking of Indels by Decomposition (TIDE) is another method with the same purpose. TIDE is based on PCR amplification, capillary sequencing and deconvolution algorithm^9^. On the other hand, Indel Detection by Amplicon Analysis (IDAA) requires a tri-primer amplification labeling and DNA capillary electrophoresis for indel detection^10^. These methods are generally cheap and quick on execution but fail to provide a comprehensive report of the gene editing outcome. Finally, Next Generation Sequencing (NGS) methods have also been broadly used to characterize gene editing efficiency^11^. NGS techniques are the optimal platforms for accurate quantification of indel size, frequency, and sequence identity determination^12^. The analysis of NGS results requires the development of bioinformatic algorithms. Some tools developed for the genome editing assessment include CRISPR-GA^13^, CRISPResso^14^, CrispRVariants^15^, CasAnalyzer^16^, cris.py^17^, CRISPResso2^18^, ampliCan^19^ and CRISPRpic^20^. Other tools are focused on the analysis of Oxford Nanopore Technologies (ONT) data, instead of Illumina-based NGS data^21^. Moreover, new analysis tools have recently been developed to cover specificity assessment related to off-targets and translocations^22^.

However, these tools have some limitations (Supplementary Table 1). For instance, quantification of different outcomes is poorly or imprecisely reported, these tools do not include simulations to help in design or benchmarking; the frequencies are based on absolute counts without taking into account amplification and sequencing biases. Preferential clustering of smaller fragments by NGS technologies has been previously shown^23^, as well as sequencing error rates^24^. Also, most of these tools lack important functionalities like reference identification, clustering, or noise subtraction.

In this context, CRISPR-A emerges as an evolution of CRISPR-GA^13^, the first NGS-based method for gene editing assessment. CRISPR-A simulated gene editing data takes into account the different repair mechanisms involved in DBS repair, and it is capable of analyzing edited bulk cell populations as well as individual clones. To address previous limitations, we have explored alignment and variant calling methods to overcome the challenges of a precise characterization.

The operation of different alignment methods, the implementation of different filters, and the correction of sequencing bias are studied to reach an accurate quantification of edition events. Development and benchmarking of CRISPR-A have been done using simulated data as well as a big variety of targeted NGS data. From CRISPR-A functionalities we highlight: (i) the development of a simulator together with the analysis pipeline, (ii) calibrated alignment method and new variant caller aware of the targeting site, (iii) batch samples analysis, (iv) nextflow pipeline implementation (https://bitbucket.org/synbiolab/crispr-a_nextflow/) that is being also implemented in nf-core, and (v) a web application with a user friendly and interactive interface (https://synbio.upf.edu/crispr-a/). In addition, it includes reports and visualizations of indels (including micro-homology patterns), substitutions and objective modification quantification, and in frame and out of frame indels report. Finally, we want to highlight the enhanced precision using Unique Molecular Identifiers (UMIs) and spike-in controls, as well as noise reduction with an empirical model based on negative control samples (mocks). Precision will be key for applications that require accurate and traceable results such as clinical CRISPR applications and environmental or industrial uses.

## Results

### Gene editing simulations provide design assessment

We developed CRISPR-A, a gene editing analyzer that can provide simulations to assess experimental design and outcomes prediction. These simulations are generated by SimGE, an R package that, for ease of use, is implemented within the CRISPR-A platform (Fig. 1A). This algorithm is useful to generate simulated data of edited reads for CRISPR analysis tools evaluation as well as for design purposes. The SimGE algorithm is based on the characterization of repair outcomes in primary T cells^25^, which are a promising cell type for therapeutic *ex vivo* genome editing. It simulates repair outcomes of CRISPR-Cas9 knockout experiments and it is able to simulate the most common variants: insertions, deletions, and substitutions, based on observed experimental data edit distributions (Fig. 1B). Same parameters and probability distributions were fitted for three other cell lines: Hek293, K562, and HCT116^26^, to make SimGE more generalizable and increase its applicability.

**Fig. 1:**
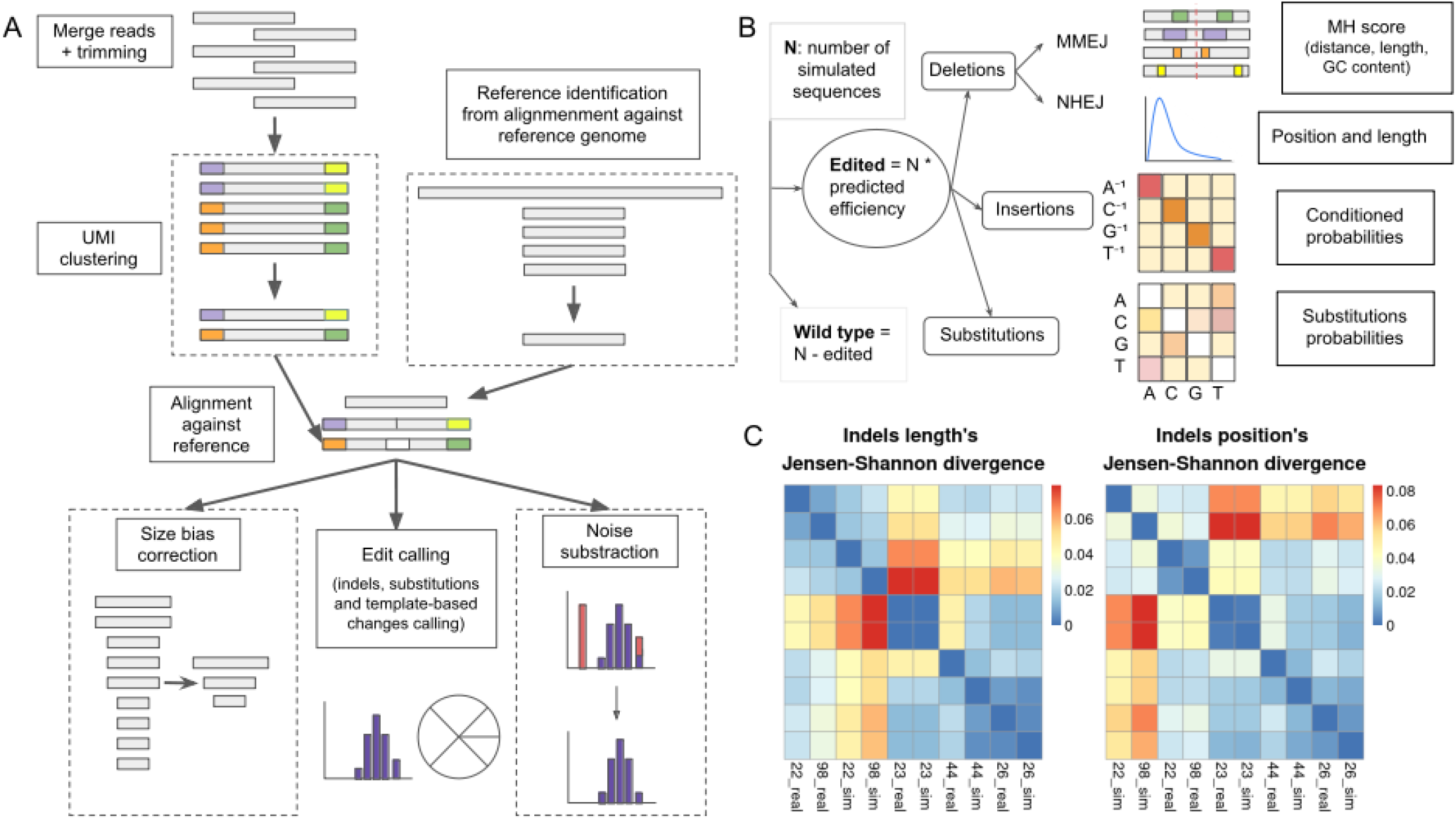
CRISPR-A capabilities for analysis and simulation of CRISPR-based experiments. **A** Schema of CRISPR-A analysis pipeline. The analysis algorithm is composed of three mandatory steps: reads pre-processing for quality assessment, reads alignment against reference amplicon, and edit calling. The processes inside dashed line squares are optional, being: UMI clustering, reference discovery, size bias correction, and noise subtraction based on an empirical model from negative control samples. **B** Squema of CRISPR-A simulation pipeline. Simulation is based on the fitting of multiple parameters that describe the distribution of edits and their characteristics and proportions. Once the number of edited sequences is determined by the protospacer predicted efficiency, other probability distributions are applied to decide the number of each kind of edit (NHEJ deletions, MMEJ deletions, insertions, and substitutions). **C** Heatmap to visualize hierarchical clustering of real samples and their simulations from validation data set. Data values are transformed to the color scale depicted on the right. Dark blue equals 0 distance and identical samples, while red is for the greatest distance value. The plot on the left shows the distances between the position of the variant and the cut site, while the plot on the right is for the size of the variants. For these plots, we have randomly selected 5 pairs of real samples from the T cells validation data set: SRR7737126, SRR7737722, SRR7737698, SRR7736744, and SRR7736723. We have labeled the samples with the two last numbers of the sample name adding real for the real sample and sim for its simulation. For instance, SRR7737722 and SRR7737698, which cluster together, are the real sample and its simulated sample for two replicates.

From the characterized data, we obtained the probability distribution of each class along with the individual DNA repair outcomes, deletions, and insertions taking into account sequence context, mutation efficiencies, and single-nucleotide variants. It was found that gamma distribution fitted best for both the insertions and substitutions whereas the beta distribution described better the size and location of the deletions. For insertions, we have observed that the majority of them, 91.79%, are 1bp (Supplementary Fig. 1), thus, SimGE predicts just insertion of 1bp size. Deletion modification sizes, as well as starting position distributions of all kinds of edits, are defined from experimental data. This ensures that the simulations are distributed in a comparable way to what we would expect after a CRISPR-Cas9 knockout experiment. After comparing the deletion and insertion size distribution from two different data sets, we observed that variants decrease quickly as the size increases (Supplementary Fig. 1), being consistent with previous observations^27^.

Among the evaluated metrics, Jensen distance showed the best clustering between real samples and replicates and, as expected, a greater separation with the unrelated samples (Supplementary Fig. 2). For this reason, this is the metric used to compare indel sizes and start positions between experimental samples and simulations. After testing the different combinations of Euclidean distances, we see that all three show a very similar clustering pattern.

To illustrate the results of the evaluation of the model, we have selected a subset of 5 experimental samples with their equivalent simulated samples. The clustering is depicted in Fig. 1C. Using Jensen distance, we see that each pair of experimental and simulated samples show a lower distance between them in all 5 different cases, compared to samples with different targets. In addition, on top of comparing the distance between the experimental sample and the simulated, we have included two experimental samples, SRR7737722 and SRR7737698, which are replicates. These two and their simulated samples show a low distance between them and a higher distance with other samples.

### CRISPR Analytics: a versatile and intuitive tool for genome editing simulation and analysis

CRISPR-A analyzes a great variety of experiments with minimal input. Single cleavage experiments, base editing (BE), prime editing (PE) or homology directed repair (HDR) can be analyzed without the need of specifying the experimental approach. Also, if the amplified genome is in the list of reference genomes, the specification of the reference sequence can be avoided, as the tool will identify which sequence must be used as an amplicon reference. Therefore, the CRISPR-A pipeline just requires NGS targeted data to compute gene editing results. Amplicon sequence, protospacer sequence, and cut site position relative to the protospacer, and template are other optional inputs.

The first step of the pipeline is the quality control of Next Generation Sequencing (NGS) raw data, which consists of merging forward and reverse reads and trimming short and low quality sequences, as well as adapter sequences. When Unique Molecular Identifiers (UMIs) are used to increase precision, reads are clustered by these sequences, and the consensus of the cluster is used in the following steps. The next step is the alignment of the reads against the amplicon sequence, which is used in the same orientation as the protospacer to achieve comparable results. After that, custom scripts have been developed for CRISPR edits calling, quantification size bias correction by spike-in controls, and mock-based noise subtraction. Finally, several interactive tables and plots are generated to visualize the results (Fig. 1A).

To standardize the analysis, CRISPR-A pipeline is implemented with Nextflow, a workflow management system that uses Docker technology for containerized computation^28^. First, NGS data can be uploaded as fastq files or it can be simulated with SimGE. Second, a reference amplicon can be directly uploaded or, in the case of an available reference genome, the reference amplicon can be identified by aligning the reads against the reference genome. Third, the protospacer sequence is also required for further standardizations in the analysis process and visualization (Supplementary Fig. 3A). SimGE, the genome editing simulator that we have assessed using five-fold cross-validation (Supplementary Fig. 3B), is also part of the pipeline. In the case of simulating edits, cell lines as well as sequence context is taken into account to determine the proportion of each outcome (Supplementary Fig. 3C). On top of the pipeline, CRISPR-A has a user-friendly and interactive web application to visualize all the results provided by the analysis. A genome browser is included to visually evaluate the performance of the alignment. Samples can be easily compared by summary tables and heatmaps. Interactive pie plots show the count of reads in all pre-processing steps, indels characterization, and edition.

Other relevant information, such as micro-homology patterns leading to deletions, substitutions, and more abundant alleles, are also shown graphically. CRISPR-A does not make assumptions during the analysis, hence, the cut site is not used for filtering as other tools use (e.g. Crispresso2, CRISPRpic, CRISPECTOR…). Nevertheless, the user has the option of refining the percentage of edition indicating the desired filtering windows iteratively once the results are shown (Supplementary Movie 1).

### CRISPR-A effectively calls indels in simulated and edited samples

Gene editing simulations obtained with SimGE were used to develop the edits calling algorithm as well as for benchmarking CRISPR-A with other tools that have similar applications. The name of each simulated edit indicates its nature and characteristics for an easier further validation of the edits classification.

Different alignment methods were compared to establish the best method to characterize CRISPR-based indels (Fig. 2A). Most of the mischaracterized alignments, for all compared alignment tools, were due to shorter and multiple indels, instead of a single deletion or insertion, which is the expected result of a DSB repair (Supplementary Fig. 4). Although all alignment methods make this kind of error, BWA-MEM^29^ is the one showing a higher abundance.

**Fig. 2:**
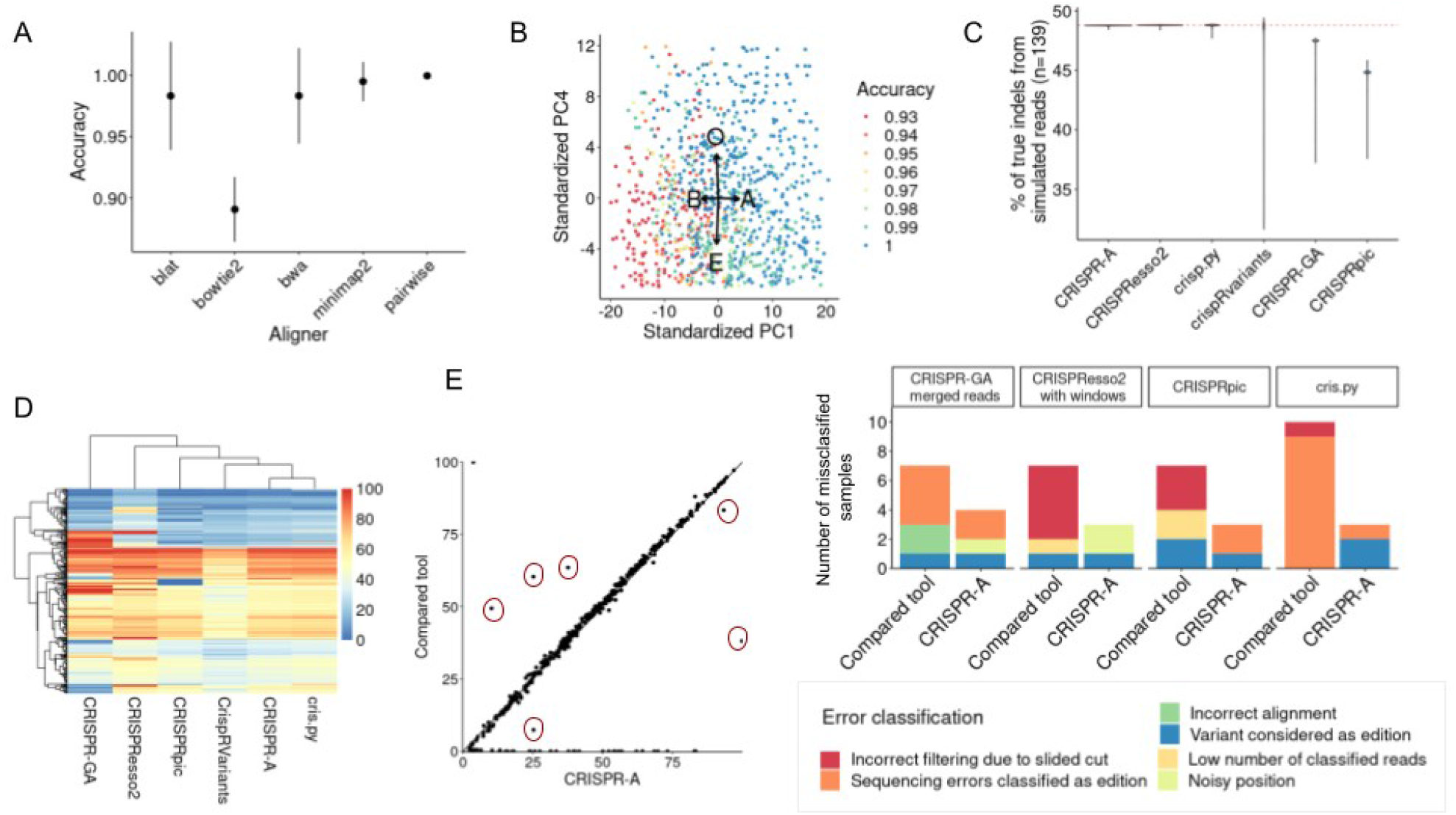
Indels characterization algorithm development and benchmark with simulated data and real data. **A** Accuracy of indels detection after aligning with different alignment tools. We have used 139 samples simulated from different reference sequences and cut site location with SimGE. The accuracy, which is the number of reads correctly classified against all classified reads, has been calculated after characterizing the indels and wt sequences of each sample after aligning the samples with 6 different methods. The mean and standard deviation of the accuracy of the different characterized samples by alignment method are represented in the plot. **B** Optimization of alignment penalty matrix. Best parameters have been determined through Monte Carlo to obtain alignments optimal for CRISPR-based indels characterization. In the PCA we can see how the four parameters of the alignment penalty matrix should be combined to achieve higher accuracy. **C** Benchmarking of indels characterization between 6 different tools. CRISPR-A is compared with 5 other tools using a simulated data set of 139 samples. All samples contain the same percentage of indels (red dashed line) and the violin plot shows us the dispersion of the reported edition percentages by each tool. **D** Reported edition of edited t-cells. 1656 unique edited genomic locations within 559 genes are characterized with 6 different tools. The percentage of edition reported by each tool for each sample is shown by the heatmap. **E** Error characterization from the most discrepant values in t-cell edited samples. CRISPR-A results are compared with the results of other tools with more distant results (example at left side; explored samples are encircled in red). Errors are classified regarding their source.

Nevertheless, we have also reported other kinds of errors. For instance, Bowtie2’s^30^ main error type is the lack of alignment, while BLAT^31^ is more prone to mislead the type of change. Pairwise alignment and minimap2^32^ show similar performance. Thus, since minimap2 is as well based on pairwise alignment and is ten-fold faster, we have focused on improving CRISPR variant alignment results with minimap2. Therefore, we have performed a Monte Carlo analysis with a random sampling of alignment parameters to increase the accuracy. Alignment and edit calling of the simulated samples were done with the parameters obtained from 950 iterations of a random selection of alignment matrix penalty values (matching score (A), mismatching score (B), gap open penalty (O), and gap extension penalty (E)). After performing a Principal Component Analysis (PCA), Fig. 2B shows a high relevance of high values in A to achieve high accuracy values, while low values of B help to obtain higher accuracies. Furthermore, high values of O and low values of E can be observed in samples characterized by high accuracy.

To evaluate the improvements brought about by the optimization of the aligner, our method is benchmarked with other currently available tools (Fig. 2C). CRISPR-A is as good as CRISPResso2 (*18*) in characterizing simulated data as both tools characterize almost all samples with a good approximation of indels percentage. CRISPRpic (*20*) has the highest mean distance to the true value (−4.03 %), followed by CRISPR-GA (−2.58 %) (*13*). CrispRvariants (*15*), even with a mean closer to the expected value, has the highest standard deviation (3.09 %).

To further explore the different tools and their performance, we compared them using targeted NGS data analysis from the primary T cells data set (*25*). When we compare the reported percentage of edition of all samples by the same 6 different tools, we see that CRISPR-A and cris.py show similar results. CrispRVariants is the one with lower reported percentages, and CRISPResso2 and CRISPR-GA have the higher amount of extreme values (Fig. 2D). To make CRISPResso2 and CRISPR-GA results more comparable with the other tools, we have given the pair-end reads assembled with PEAR to CRISPR-GA, and the protospacer to filter by edition window to CRISPResso2. Then, we see a reduction of extreme values in CRISPResso2, and a reduction of dispersed values in CRISPR-GA (Supplementary Fig. 5). After that, we also compared the edition percentage reported by all these tools against the indels percentage given by CRISPR-A. CRISPRpic shows a high number of samples with null edition percentage, while other samples are giving edition values for these same samples. CrispRVariants shows, in general, lower reported percentages compared with the other tools, and a higher dispersion for indels with higher edition (Supplementary Fig. 5). In addition, we explored more in detail the samples with the highest difference of reported percentage of edition compared with our tool (Table S2). After comparing in detail these samples we can conclude that CRISPRpic and CRISPResso2 tend to report single nucleotide polymorphisms (SNPs) as edits and remove true indels produced by CRISPR systems when the cut site is slided, while CRISPR-A and cris.py tend to report error prone sequence regions as edits (Fig. 2E). We have investigated the origin of these errors done by CRISPR-A to improve our tool. There are 6 samples in which CRISPR-A has been considered to be giving an incorrect edition percentage report. There is one sample, SRR7736645, whose amplicon sequence does not coincide with the one of the reference genome. It has an heterozygous variant, an inversion in one allele, and an insertion in the same position in the other allele (Supplementary Fig. 6). Since the mutation is not close to the cleavage site, in this case, the tools that apply a filtering window are the ones giving a better approximation of the edition.

Samples SRR7736865, SRR7737875, and SRR7737569 have been analyzed with a sequence shorter than the amplicon sequence and an incorrect protospacer. For this reason, all samples are reporting a lower percentage of edition. Once the reference sequence is corrected, using CRISPR-A amplicon sequence discovery function, CRISPR-A shows a perfect edition profile (Supplementary Fig. 7). Finally, samples SRR7736598 and SRR7736646 have a polyadenine sequence of length 14 that is poorly sequenced (Supplementary Fig. 8). For this reason, the tools with filtering windows have a better estimation of the edition.

### Improved discovery and characterization of template-based alleles or objective modifications

CRISPR-A can be used to look for any kind of genetic mutation if a sequence with the change and its surrounding homology sequences are given. This allows reporting of HDR, PE, BE, or even microhomology-mediated end-joining (MMEJ) or NHEJ alleles of interest. Of the explored tools for indels characterization, CRISPResso2, CRISPR-GA, and CRISPR-A are the only tools capable of analyzing HDR events. These tools report values close to the expected percentage of template-based or objective modifications when simulated data is analyzed (Fig. 3A). We added the same amount of reads with certain template-based modifications to the simulated data obtained with SimGE. Even if the amount of sequences with modifications is the same for each sample, the characteristic (size, position, and nature) of each modification is different to have a comprehensive evaluation of the performance. The mean difference to the real percentage of sequences with the objective modification (1.09%) is 2-fold higher in CRISPResso2 than in CRISPR-A, while CRISPR-GA shows the closest mean distance to this value. Even so, there are some samples with a higher objective modification count, corresponding to indels that lead to the same modification as the template-based change. In general, CRISPR-A distribution is closer to expected, centered on the percentage of template based added reads, and with low dispersion.

**Fig. 3:**
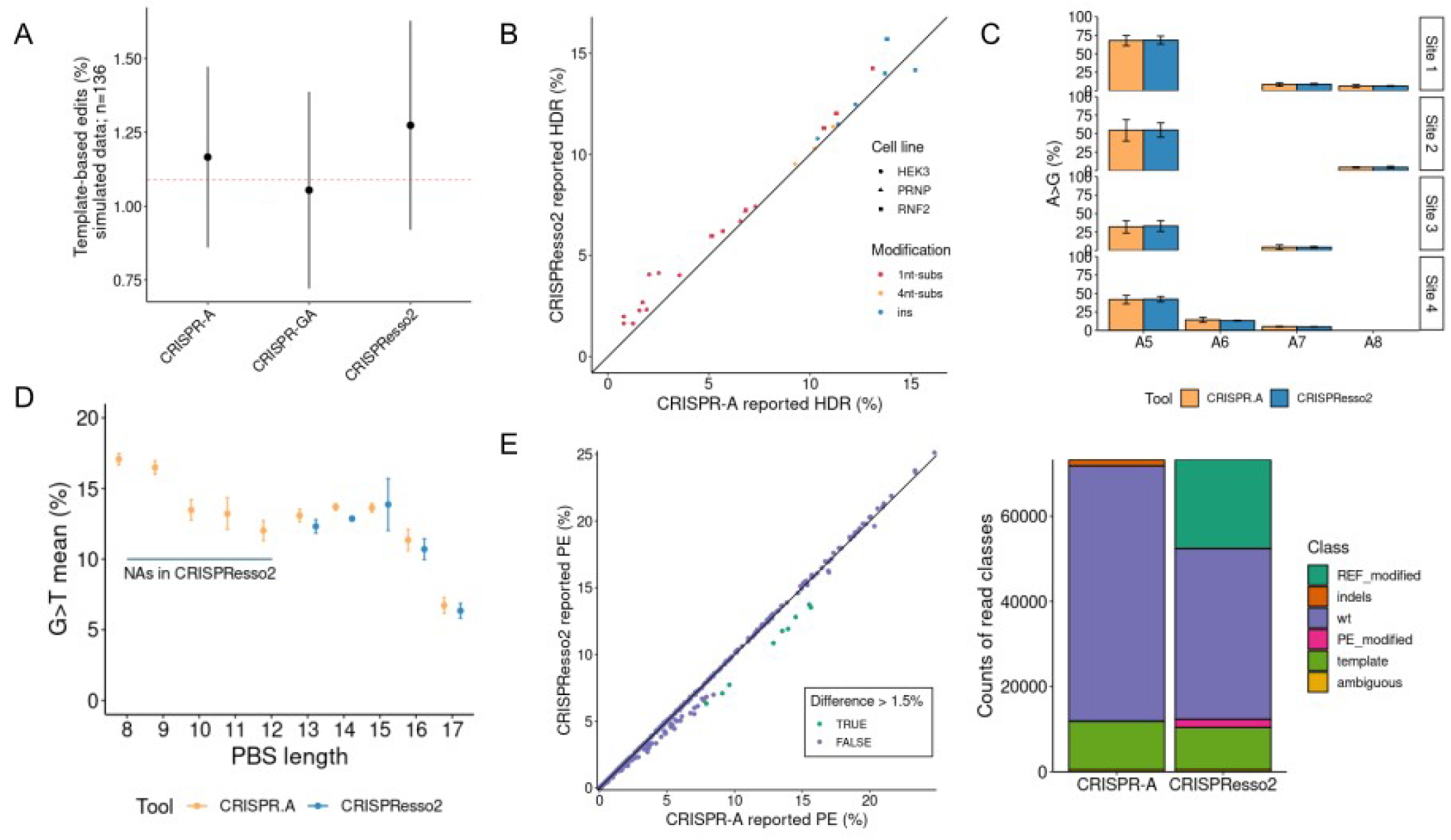
Template-based edits and substitutions characterization benchmark with simulated data and real data. **A** Benchmarking of template-based edition characterization between 3 different tools. CRISPR-A is compared with 2 other tools using a simulated data set of 136 samples. All samples contain the same percentage of template-based reads (red dashed line). We can observe the mean template-based percentage among all characterized reads in each sample and the standard deviation reported by each tool: CRISPR-A, CRISPR-GA, and CRISPResso2. **B** Characterization of different modifications obtained by HDR. CRISPR-A and CRISPResso2 are used to characterize the result of 27 experiments with different modifications led by HDR: substitutions of one nucleotide by another, substitutions of 4 nucleotides for other 4 nucleotides, and insertions of 3 nucleotides. The modifications are found in 3 different targets. **C** Characterization of substitutions done by BE. In this case, instead of looking for the overall percentage of edition we looked for the information of substitutions by position given by two different tools: CRISPR-A and CRISPResso2. There are 4 different targets and 3 replicates for each target. **D** Characterization of PE substitutions. We have compared a total of 29 samples of PE edition of FANCF gene with templates with different PBS lengths. With CRISPResso2 analysis, the examples with a PBS shorter than 13 base pairs end up with error, and results are not obtained. **E** Extended analysis of PE substitutions. 23 different samples edited in 5 different targets (HEK3, EMX1, FANCF, RNF2, and HEK4) are analyzed with CRISPR-A and CRISPResso2. In the plot on the left side, we can see the comparison of the reported PE percentage by each of the two tools. HEK3 edited samples have a higher PE edition reported by CRISPR-A than with CRISPResso2 (in orange). In the right part of the figure, we found the count of reads classified in each class. CRISPResso2 reports a high number of PE with modifications or reference with modifications due to a SNP (rs1572905; G>A).

CRISPResso2 shows a higher dispersed distribution, while CRISPR-GA tends to underestimate the objective modification percentage.

In addition to simulated data, we have analyzed targeted NGS data of HDR, BE, and PE experiments. The targeted NGS analyzed data of an HDR experiment^33^ shows that CRISPResso2 is reporting slightly higher but significant HDR percentages than CRISPR-A in most of the cases (p<0.00005) (Fig. 3B) but it is difficult to explain the origin of these differences. However, what we can observe after exploring in detail 6 of the HDR analyzed samples (Table S3) is that CRISPResso2 is classifying more reads as template-based than CRISPR-A, while CRISPR-A, in most of the cases, is classifying more reads as reads containing indels than CRISPResso2 does. The samples with higher differences, almost 2-fold change, have a 42% of the reads classified as ambiguous by CRISPResso2, probably due to a substitution present in two thirds of the reads.

Base edited samples ^34^ can also be analyzed as objective modifications by CRISPR-A if the percentage of a specific change in a particular position wants to be explored. In addition, CRISPR-A always searches substitutions along the whole amplicon sequence that can be used when all modifications at the edition windows are of interest. CRISPResso2 and CRISPR-A give comparable results of substitution quantification in base editing experiments, using the default substitution search along the whole amplicon sequence done by CRISPR-A and indicating base editors as the editing tool to CRISPResso2 (Fig. 3C). Prime editing (PE) is another case in which a template, the 3’ extension that is composed of Primer Binding Site (PBS) and reverse-template, is used to achieve an objective modification. We have analyzed FANCF gene modifications made with PE^33^ and, even though results are similar when PBS is longer than 13 nucleotides, CRISPResso2 is not able to give results for PBS shorter than 13 nucleotides for this target, due to misleading alignment between 3’ extension and amplicon sequence (Fig. 3D). We have extended these analyzes adding more samples from the same project and we observe that CRISPR-A gives higher editing percentages of template-based edition in several cases (Fig. 3E). We have explored more in depth the cases where the difference was higher than 1.5%. This difference is explained by a high number of reads classified as reads with reference modifications or prime editing modifications by CRISPResso2. The sum of all these ambiguously classified reads is almost equal to the sum of the missing wild type, indels, or PE reads reported by CRISPR-A. All samples with these differences have the same target, and the undetermination is due to the fact that there is a SNP (rs1572905) in heterozygosity.

### Characterization of cell line dependent and independent editing outcomes with CRISPR-A

Three different cell lines: HEK293, K562, and HCT116 were analyzed with the CRISPR-A analysis pipeline. CRISPR-A has enough precision to find relevant features and outcomes in different cell lines. The distance between the percentage of microhomology mediated end-joining deletions of samples with the same target was calculated and the mean of all these distances was used to reduce the information of the 96 different targets to a single one. After this, the replicates of each cell line cluster together (Fig. 4A). In addition, we have done a differential expression analysis and we have found that *POLQ* and *RBBP8*, also known as *CtlP*, are significantly more expressed in HCT116 than in K562, which are the cell lines with the major and minor ratios of MMEJ compared with NHEJ, respectively (Fig. 4B). POLQ is essential for MMEJ in all mammalian species^35^ and CtlP promotes MMEJ repair^36^. We also observe differences in the fraction of insertions among all indels, being HCT116 the cell line with the highest proportion of indels (Fig. 4C). When we do an overall visualization of the percentage of indels observed in each sample, we also see different patterns among cell lines, with HCT116, a repair-deficient cell line, and HEK293 showing a higher percentage of edition than K562 (Fig. 4D).

**Fig. 4:**
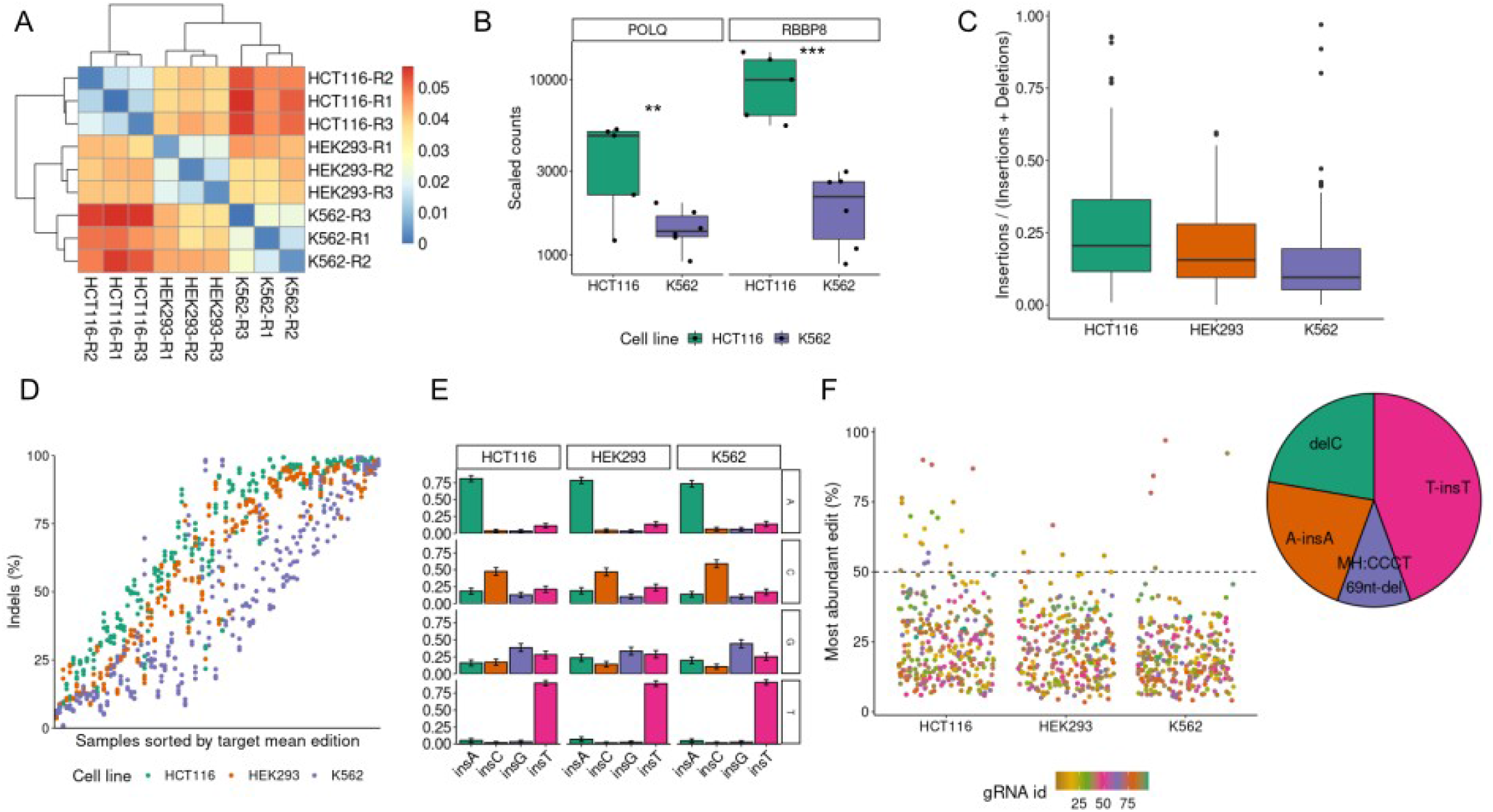
HCT116, HEK-293, and K562 edited in 96 different targets analyzed by CRISPR-A. **A** MMEJ deletions ratio among all deletions. We have calculated the euclidean distance of the ratio of MMEJ deletions among all characterized deletions. The heatmap represents the mean between the distances calculated for each of the 96 different targets. The three replicates of each cell line are clustered together showing that the MMEJ ratio is a feature that depends on the cell line. **B** Insertions against total edits. The proportion of insertions among all reported edits in three different cell lines. **C** Percentage of editions. Indels among all characterized reads in 96 different targets with 3 replicates by each target in three different cell lines. **D** Insertions considering the nucleotide upstream of the cut site. Percentages of inserted nucleotides in the function of the free nucleotide in three different cell lines. **E** Variant diversity by samples. On the left is the percentage of the most abundant variant among all kinds of edits. In the right the distributions of variants from all the variants that are above 50% (dashed line in left plot), these are the variants with higher abundance.

There are other CRISPR-based editing outcomes that instead of being characteristic of each cell line are common for all of them. This is the case of insertions of a single nucleotide at the cutting site, which shows a strong prevalence of insertion homology when there is thymine or adenine. This is not as frequent when the free nucleotide is a cytosine, and even less when it is guanine (Fig. 4E). Another common feature is the diversity of variants by targets. We have explored the targets that have an edition outcome that represents more than half of all different edits in a certain sample. Of the 96 gRNAs, there are 9 different gRNAs which lead to outcomes with low diversity of editing genotypes. We can see certain strong patterns on these targets: free thymine or adenine at the 3’ nucleotide upstream of the cutting site that lead to insertions of the same nucleotide, a free cytosine at the same place that leads to its loss, and strong mico-homology patterns that lead to a long deletion (Fig. 4F).

### Three different approaches to increase precise allele counts determination

We have implemented three different approaches to increase edits quantification accuracy: (i) synthetic molecules of known size and quantity (spike-in controls) to model size biases, (ii) UMIs to remove PCR duplicates, and (iii) an empirical model based on mock samples to denoise the treated ones.

Synthetic molecules with known deletions and insertions of different sizes in relation to the reference sequence were analyzed together with edited C2C12 cells genomes. The differences in the final count of the spikes increased together with the number of Polymerase Chain Reaction (PCR) cycles, and the number of initial molecules is not as relevant as the number of cycles (Supplementary Fig. 9). After observing the linear correlation between the size of the spikes and their relative count, we have fitted a linear model to transform the indels count depending on its difference in relation to the reference amplicon (Fig. 5A). When we compare the original count with the updated counts after size-based correction, we see that the distribution of indels is flatter (Fig. 5B).

**Fig. 5:**
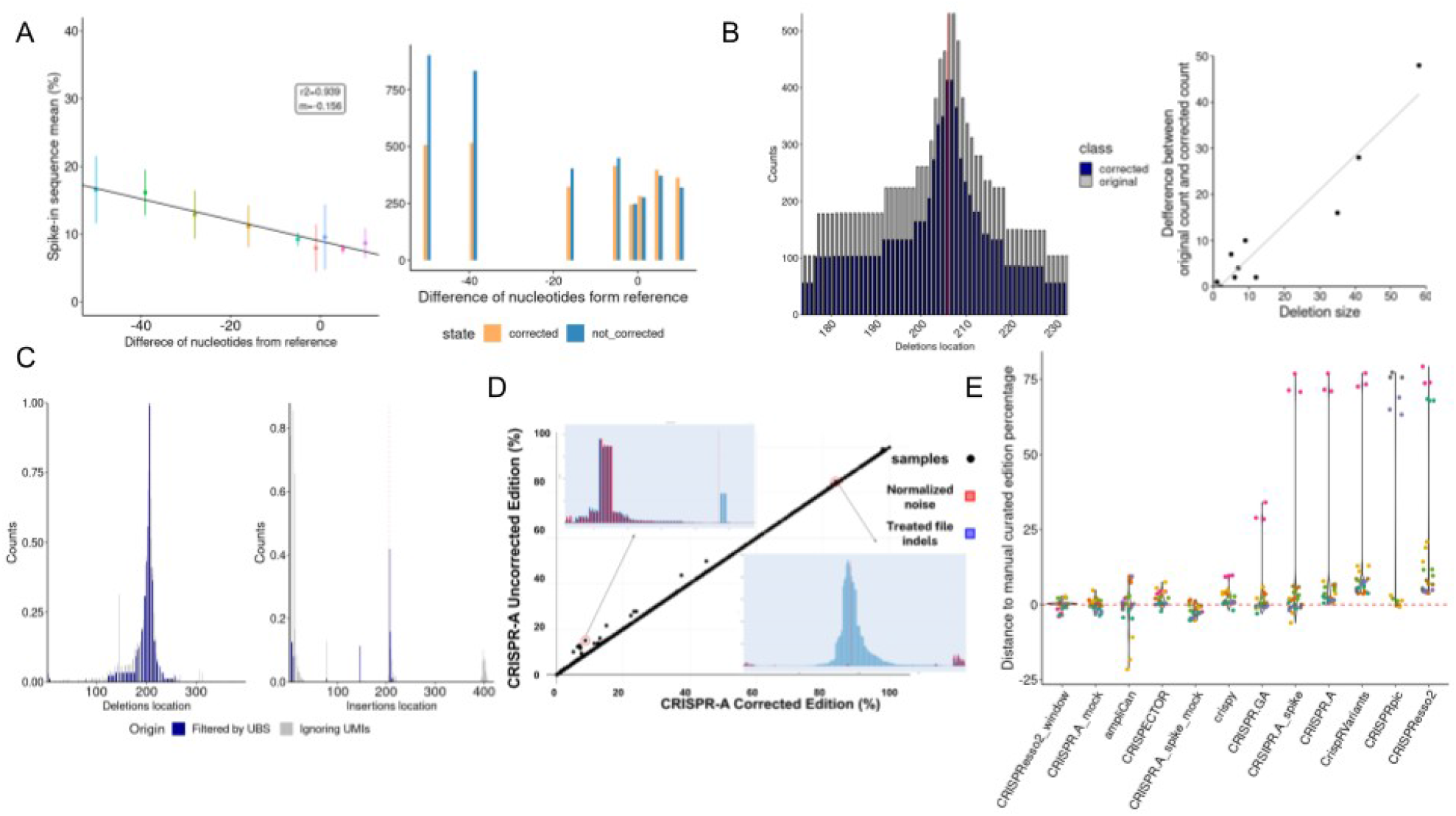
Enhanced precision with spikes, UMIs, and mock characterization. **A** Spikes count in the Illumina experiment. Count of spike-in synthetic sequences with different deletion sizes. From each spike, the same number of molecules were added to the edited samples. On the left, linear regression of spike-in sequences mean percentages among all spike-in sequences at 30 cycles of amplification and a low number of molecules. At the right, count of spikes in the original sample and after correction by spikes. **B** Size bias correction using the spike-in model. On the left, edited sample deletions distribution by position corrected by spikes (blue) against the original distribution (gray). At the bottom right, count difference between original and corrected in function for deletion size of sample deletions distribution shown at left. **C** Noise reduction by UMIs cluster filtering. Standardized distribution of deletions (left) and insertions (right) without taking into account UMIs (gray) and after clustering by UMIs with a minimal identity of 0.95 and filtering by UMI bin size (UBS) > 50 and < 130 (blue) in Lama2 target. The Red dashed line corresponds to the cut site position. **D** Mock-based noise correction. Samples with less percentage of edition tend to have a higher correction since the noise represents a higher proportion of the indel reads. The two plots comparing treated and mock files show that this subtraction is always specific, regardless of the edition percentage. **E** Difference between the edition percentage reported by 8 tools and the 4 options of CRISPR-A and the manually curated percentage of 30 samples (10 different targets and 3 replicates for each).

Using UMIs we can cluster several reads that come from the same original molecule and see if the indels and substitutions found are errors or are real changes produced before the sequencing process. We have used two different amplicons (TRAC and Lama2) to explore the percentage of reported indels equal to the consensus of the cluster. The explored clusters were generated using different minimum allowed sequence identities. The identity is the number of alignment columns containing matched residues between the length of columns after alignment; we explored 80, 85, 90, 95, and 99 minimum allowed identity within the clusters. We saw that smaller clusters tend to show less percentage of reads supporting the consensus sequence, while with clusters that have an identity lower than 90% the dispersion is maintained since bigger clusters are achieved (Supplementary Fig. 10A).

One example, where the effect of using UMIs, clustered by an identity of 95, to filter data can be seen is shown in Fig. 5C. The sequenced target has a high sequencing complexity that leads to imprecise sequencing. In gray, we can see indels normalized counts before applying UMIs clustering and cluster filtering. After filtering by UMI bin size (UBS) higher than 50, removing the sequences with the same origin but contained in several clusters, and by UBS lower than 130, removing sequences with different origins in the same cluster, we see that noisy positions are filtered. Removal of noise using the same approach can also be seen using other targets as TRAC (Supplementary Fig. 10B).

Lastly, CRISPR-A uses an empirical approach to address the misled classification of errors as editing events, requiring a negative control (mock file) to denoise treated samples (Fig. 5D). This step of the analysis is optional and flexible, allowing combinations of samples to be corrected with the same or different mock files. The model compares the indels occurring in the mock and in the treated file separating by indel type (insertion or deletion), size in bp, and position relative to the cut site to avoid bias since significant differences in these features were observed between the indels observed in the mock and in the treated files present in a data set with three different cell lines (HEK293, K562, and HCT116) that had negative controls for the three biological replicates of each gRNA (Supplementary Fig. 11 A, B, and C).

Finally, we have compared the different versions of CRISPR-A (using or not a negative control, and correcting or not size bias by spike-in control) and all other tools explored in the benchmarking with a manually curated data set (Fig. 5E). The use of a negative control sample when running CRISPR-A allows the obtention of results with a lower difference to the manually curated value. CRISPResso2 has a high dependency on the quantification window determined by the cutting site. CRISPRpic is the tool with less accuracy detected since it only accurately characterized 18 of the 30 analyzed samples and is far from the correct edition percentage in other 6 samples. CRISPR-GA, cris.py, CripRVariants, and CRISPECTOR show in general values higher than the manual curated value. AmpliCan has outliers on both sides. The tool with a mean closer to 0 is CRISPR-A denoised using a negative control.

### CRISPR-A empiric model removes more noise than other approaches

The fact that a model subtracts more or fewer indels does not necessarily mean that it removes more noise, since it may be masking bona fide indels. Therefore, to establish a framework on which we can compare our model we looked for how many indels with the same characteristics (class, position, and size), we could find at the same time in the mock and treated samples from the data set that had the negative controls (Supplementary Fig. 12A). What we observed is that the representativeness of the mock files, although quite high near the cut site, is not total in the rest of the sequence, i.e, that there are indels introduced by the noise in the mock files that do not have an equivalent in the treated ones. Moreover, when calculating the proportion of treated file indels represented in their mock files (Supplementary Fig. 12B) we found that it is much more likely that the reported indels far from the cutting site are noise since they are proportionally more represented. Knowing this, we calculated the subtraction efficiency of our model (Supplementary Fig. 12C), obtaining that, of the indels represented, those that are in higher proportion (far from the cut site) are almost completely subtracted while those that are less proportionally represented (close to the cut site) are less subtracted. It is also observed that after normalizing by aligned read depth, these characteristics improve (Supplementary Fig. 12D) suggesting that normalization and proportional subtraction generate an efficient and sensitive model by minimizing the possibility of masking bona fide edits that are represented by chance.

After characterizing the efficiency and sensitivity of our model, we could do benchmarking against other denoise approaches. This would let us know if this novel approach improved somehow the state-of-the-art. Among all the tools presented in the introduction, only ampliCan ^19^ and CRISPECTOR^22^ apply some kind of noise correction model. To benchmark the CRISPR-A model, we compared its performance with the previously characterized data set.

AmpliCan evaluates position by position the amount of noise, and if it exceeds a certain threshold, it considers all indels in that position as noisy, regardless of the class. Meanwhile, CRISPECTOR uses a Bayesian model with configurable priors that evaluate positions as noisy or not, stratifying by indel type, and position.

The results were surprising, as the subtraction applied by ampliCan had zero effect on the reported editing percentage. In the case of CRISPECTOR, a correction, which is a change in the editing percentage using a mock to denoise compared without using it, was observed, although very modest, since only about 10% of the files underwent a correction. For CRISPR-A this percentage went up to 82% (Supplementary Fig. 13A).

To better understand where these differences may be occurring, we explored the correction of some of these samples. It should be noted that this comparison was difficult to evaluate as CRISPECTOR only provides the percentage of reads edited, without specifying the position or any other information that allows us to track them.

For instance, sample SRR3700075 has a repetitive sequence region at the beginning of the sequence (*CTG CTG CTG CTG CAG CAG CAG*) that produces a peak of indels, probably due to noise. This peak partially masks the peak next to the cut site. CRISPR-A effectively corrects this region since it is equally represented in the mock, while CRISPECTOR shows a correction close to zero.

An opposite case is SRR3696877, where we observe a greater correction of CRISPECTOR than the one applied by CRISPR-A. As can be seen in Supplementary Fig. 13B, besides a few noisy insertions at the extremes, there’s not much subtraction going on. Some deletions next to the cut site were tagged as noisy, but not subtracted. It is difficult to assess if those deletions correspond to the extra correction that CRISPECTOR performed, but CRISPR-A limited this subtraction since, in the mock file on those positions, the observed normalized frequency was lower than the treated, avoiding masking.

Finally, it is worth mentioning that there are samples that CRISPECTOR does not analyze due to an imbalance between mock and treated samples. Since we saw no correlation between depth reads and the representativity (Supplementary Fig. 14), we concluded that the subtraction can still be performed. That’s why CRISPR-A does denoise the samples while notifying the user that the depth is lower than the arbitrary threshold of 300 reads, as in SRR3698365.

## Discussion

NGS is the method that enables the identification of all different outcomes led by genome editing tools. There are different online and command line available tools to decipher the percentage of edits achieved in genome editing experiments. Even so, most of these tools do not retrieve all possible kinds of edition events and are not flexible enough to cover the whole diversity of genome editing tools. Moreover, none of them include simulation to help in the design or analysis performance evaluation. Furthermore, alignment, amplification and sequencing errors have not been previously taken into account systematically to achieve a precise estimation of CRISPR-based experiments results.

Here we described CRISPR-A, a tool that can be used to simulate gene editing data as well as analyze the efficiency and outcomes profile of any kind of CRISPR-based experiment. Our tool has achieved a higher performancer to those of current applications, with additional versatility, and with a dynamic oriented reporting that enables its use by a wide scope of users. In CRISPR-A analysis, there are no biases due to assumptions related to the performance of the used technology since the analyzes are agnostic to the used gene editing tool and the denoising process is done using an empiric model. This makes the tool suitable for the fast development of the genome editing field. The novelty of the applied methods such as the use of UMIs and spike-in controls for quantification correction for an enhancement in precision is unique among the current CRISPR-based analysis tools and it may facilitate use in situations where accuracy is paramount (i.e: clinical). We have also improved the use of mock samples in the correction of confounding errors.

We have coupled the analysis pipeline with a simulator that has been useful to benchmark gene editing tools. The probability distributions of the algorithm were fitted with 4 different cell lines to capture the peculiarities of each one of them, which makes it more generalizable. After exploring different distance metrics, we determined that Jensen-Shannon divergence is the best measure of these probability distributions similarity. We succeeded, since the distance within samples with the same target, simulated and real, had a lower distance than between different sample targets. On the side of the analysis, alignment matrix penalty values were tuned to obtain a better edit calling. When current tools were benchmarked with simulated data, even without using spikes or mock correction, only CRISPResso2 approached CRISPR-A accuracy. While with real data, these two tools can only be compared if CRISPResso2 uses a filtering window based on the cut site inferred from the protospacer sequence. This implies a poorer edition report of CRISPResso2, compared with CRISPR-A, in the samples where the cleavage site is slided. Even so, window filtering is convenient to remove noise, for that reason we have implemented other approaches with this aim like an empirical model based on mock samples that is useful to eliminate the noise produced from multiple sources, such as sequencing errors, misalignments, natural variants, etc. CRISPR-A with mock correction is the most accurate tool. In addition, after the analysis CRISPR-A allows filtering by windows interactively; this is helpful, when a mock sample is not available, to remove alignment errors in the primer binding site that can be observed in the extremes of the amplicon. In addition to double strand break characterization, CRISPR-A looks for any kind of genetic mutation when a sequence with the change and its surrounding homology sequences are given. Even when an objective modification is not given, CRISPR-A reports all found substitutions along the amplicon. Successful results with different kinds of data and approaches were achieved. One limitation of the agnostic search of substitutions is that the report is not given decombuled by substitutions in the same read, it just gives an overall percentage by position. Meanwhile, the obtained results of objective modifications show an improved performance compared with current tools.

We have used CRISPR-A to characterize gene editing outcomes in three relevant cell lines since our tool is precise enough to discover relevant features. We have found certain patterns associated with low diversity outcomes: free thymine or adenine at the 3’ nucleotide upstream of the cut site that leads to insertions of the same nucleotide, a free cytosine at the same place that leads to its loss, and strong mico-homology patterns that lead to long deletions. These kinds of patterns that lead to a certain repair resolution with low variability are of high interest when a certain consequence of the cut has to be achieved. For this reason, CRISPR-A shows the edited variants with higher prevalence in analyzed samples.

Precision has been enhanced in CRISPR-A through three different approaches. The observed inverse correlation between amplicon size and over amplification, has been used to correct size biases and obtain flatter distributions of indels. We also removed indels in noisy positions when the consensus of clusterized sequences by UMI, are used after filtering by UBS. Finally, an empirical model based on mock samples has been implemented to denoise the treated ones exceeding the tools that implement similar methods.

While CRISPR-A provides a high performance platform for gene editing analysis, it will continue evolving to incorporate future improvements. There is still room to explore UMIs and spike-in correction methods more in-depth. At that moment, the analysis pipeline that clusters UMIs is independent of the nature of the sequencing technology used. In the case of spike-in controls, a more complex model simulating PCR amplification could be explored and developed. Another area of improvement is the noise subtraction model since it requires a negative control which is not always available. To solve this, a mock-independent model that takes into account the trends that we have observed in noise, together with generally described noise signatures such as homopolymeric regions or high/low GC content^37^, could be implemented to follow CRISPR-A minimal assumption philosophy.

## Methods

### Simulations algorithm development

SimGE is built taking into account the different layers of classes and their proportions. The proportion of edited and not edited sequences can be determined by two different sgRNA efficiency predictors, Moreno-Mateos^38^ and Doench 2016^39^ scores, which give the most reliable on-target activity prediction^40^. Both models give a group of weights that are assigned as descriptors of the gRNA sequence in order to define the efficiency as a value that falls between 0 and 1, being 1 the most efficient gRNA. Depending on the experimental design, one model or another suit better the data: for guides expressed in cells from exogenous promoters, like U6, Doench 2016 scores are recommended, but for guides transcribed *in vitro* from T7 RNA polymerase promoter, using Moreno-Mateos scores is a better option.

Deletions and insertions location distributions are determined by the fitted distributions. The same happens with deletions sizes. Insertions, by default, have a size of 1 nucleotide. The inserted nucleotide is determined by a conditioned probability, taking into account the nucleotide upstream to the cut site, since insertion homology is a feature with high prevalence^41^.

Furthermore, the sequence context is more relevant in microhomology mediated end-joining (MMEJ) deletions than in non-homologous end-joining (NHEJ) deletions, since for MMEJ deletion an homological pattern on both sides of the cut site is required. For the NHEJ ones, the specific nucleotides are removed based on the probability density function of the sizes and the probability distribution of the positions relative to the cut site. To determine the proportion of each MMEJ deletion and NHEJ:MMEJ ratio, we calculate the microhomology (MH) score based on the MH pattern and the length of the deletion, similarly as done by Bae et al. ^42^.

### Simulations evaluation

With the purpose of being able to compare the different sample sizes and the positions of the indels, we needed to define a distance metric. When clustering the observations into groups, we computed the distance between each pair of observations (Table S4), giving an idea about the dissimilarity among the observations. The explored distance metrics were: Euclidean mean, Euclidean median, Kullback-leibler, and Jensen. Euclidean distance is defined as the length of a line segment between two points in Euclidean space. If columns have values with differing scales, like the indels for different samples, it is common to normalize or standardize the numerical values across all columns prior to calculating the Euclidean distance. Otherwise, columns that have large values will dominate the distance measure. Euclidean distance is calculated as the square root of the sum of the squared differences between the two. Using the median instead of the mean and a more robust dissimilarity metric is much less sensitive to outliers. Median and quantiles are not suitable to describe the distribution if we have few categories, being the standard deviation a better fit. Previously, mutual information, entropy, and Kullback-Leibler (KL) distance have been used to study different parameters^43^. Hu and Hong use KL with the goal of studying the changes in isoenzyme expression, being the entropy, a measure to quantify complexity, predictability, and progression patterns. Jensen-Shannon (JS) divergence and distance have been used to explore probability distributions, being considered the symmetric and smoother version of KL divergence. Using the Jensen-Shannon and Kullback-Leibler metrics, we aimed to analyze both size and position of indels between samples.

We used five-fold cross-validation to train SimGE. We split the T cell data^25^, which includes 1.521 unique cut sites, into five-folds and trained SimGE on four of the five folds (Supplementary Fig. 3B). The same process was performed for the other three different cell lines, including 96 unique cut sites and three replicates^26^. Afterward, we tested the performance on the fifth fold, generating the simulated sequences with the same target and gRNA as the samples that belong to the fifth fold, in order to calculate the distance between these. The final validation, with the mean parameters of the different training interactions, was performed on a testing data set that was not used in the training. Validation was done with samples that had never taken place in the training process. Jensen distance is used to compare the characterization of real samples and simulated samples since this is the explored distance that differentiates better replicates among samples

### CRISPR-A gene editing analysis pipeline

When paired-end reads are used, the merge of the reads is done with PEAR^44^. After that, FastQC^45^ and Cutadapt^46^ are used for the detection and removal of adapters. Then, quality filtering is done with fastq_quality_trimmer from fastx-toolkit^47^ with a quality threshold of 20 and a minimum length of 80. An adapted version of extract_umis.py script from pipeline_umi_amplicon pipeline (distributed by ONT https://github.com/nanoporetech/pipeline-umi-amplicon) is used to get UMI sequences from the reads, when the three PCRs experimental protocol is applied. Then vsearch^48^ is used to cluster UMI sequences. UMIs are polished using minimap2^32^ and racon^49^ and consensus sequences are obtained using minialign (https://github.com/ocxtal/minialign) and medaka (https://github.com/nanoporetech/medaka).

If instead of the amplicon reference sequence the genome reference is used, reads are aligned using BWA-MEM^29^. After that, samtools^50^, bedtools^51^, and custom scripts are used to get the reference amplicon sequence. This process is based on alignment coverage and length of the region with aligned reads above it. Before aligning reads against the amplicon reference sequence with minimap2^32^ using the following parameters: -A 29 -B 17 -O 25 -E 2, the amplicon reference sequence is placed in the same orientation as the gRNA to get standardized reports for all analyzed samples. Instead of using minimap, other aligners such as BWA-MEM^29^ and Bowtie2^30^ can be used.

Variant calling or edit calling is done based on the cigar sequence and using custom scripts. Deletion’s position and extension are determined by being aware of the nature of the cut and repair mechanisms. For that reason, the deletions considered are not truncated and when different possibilities arise, the expression of deletions corresponding to those that are above the cut site are chosen. Substitutions by positions are also reported using pileup from samtools^50^, and in case of the existence of an objective modification, we look for it by aligning the reference against the modified reference and counting the cigars with the same pattern. When there are no deletions or indels in the cigar, substitutions are searched after doing pairwise alignment to get the position, size, and nature of the substitution.

Also in the variant calling process, spike experimental data is used to correct size biases of deletions and insertions. The proportion of reads is removed (for deletions) or added (for insertions) taking into account the linear regression observed when spikes of different sizes and at the same initial amount were counted. The transformation is done using the following formula, where *m* is the slope of the linear regression and *n* is the origin:

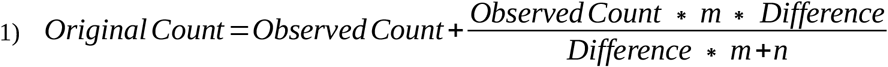

If the amplified region of the treated samples is also amplified without treatment (mock), this negative control is used to remove noise.

### CRISPR-A noise subtraction pipeline

To apply the subtraction, the mock file is analyzed in an agnostic manner, following the same path as treated files, in an independent manner. The idea behind it is that the complexity of sources that can lead to technical noise should be represented on a control that has undergone the same process with the same sequence. Thus, after performing the edit calling, the mock samples are paired with their respective treated samples.

The first step is a recovery of truncated events, in which reads classified as truncated in the treated file are compared against the mock. This allows to recover real edits that could not be classified due to a noise event occurring in the same read. Right after aligned read depth normalization occurs, we obtain a normalization ratio using the following equation:

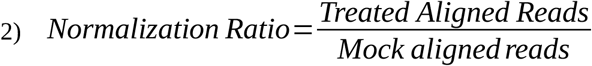

This ratio is then used to multiply the number of noise events found on the mock, gathered by length position and indel class, to avoid biases. Note that this also occurs when there are more aligned reads on the mock than on the indels, but instead of multiplying, it divides the indels to avoid over subtraction. After normalization, a simple subtraction is performed based on the number of indels observed in the control, being subtracted from the treated file only those indels that match in position, length, and edit class (insertion or deletion). After this process, if there are still reads in the treated file that meet these conditions but that have not been subtracted because they occur proportionally more frequently in the treated file, they are tagged as noisy reads differentiating them regarding their relative frequency, i.e, how big was the difference in counts between them. The reads subtracted are then classified as wild type and, for that, indels and wild type percentages are recalculated. All these processes are based on R 3.6. and included as a nextflow pipeline.

### Alignment and gene editing analysis tools benchmarking with simulated and real data

SimGE was used to simulate more than one hundred sequenced edited targets. The coordinates of the sequences were randomly selected from the human genome (hg38) with bedtools. Samtools was used to get the fasta sequences and a custom script was used to select a SpCas9 possible cut site at least 35 nucleotides away from both extremes of the sequence. After converting the fastas with the simulated edits into fastqs with SeqIO from Biopython, reads were aligned against reference amplicon sequences using BWA-MEM, BLAT, minimap2, and pairwise alignments to compare their performance. The accuracy of the alignment of each sample by each aligner was computed dividing the number of reads correctly classified by the total amount of reads. Pairwise alignment was the slowest option and minimap2, was the second best aligner. We explored the performance of minimap2 tuning the matching score, mismatching penalty, gap open penalty, and gap extension penalty. To achieve it, we have done a random sampling of parameter combinations and we have analyzed the alignments to calculate again the accuracy for all samples. From PCA analysis we can see the relevance and the space of combinations that move us to more accurate results (Fig. 2B).

These simulated data sets were also analyzed with CRISPR-A after updating the alignment parameters and other popular tools (CRIPRResso2, CRISPR-GA, crispr.py, crispRVariants, and CRIPRpic. CRISPREsso2 (version 2.2.5) was run with docker and using default parameters.

CRISPR-GA was run through its web application. R 3.6.0 was used to analyze data with CrispRVariants (version 1.14.0). In this case, we have just used the function *indelPercent* to get the percentage from reads aligned with BWA-MEM as in their usage examples. We have used an environment of python 2.7 to run crispr.py v2 and CRISPRpic.

We also analyzed the primary T cells data set^25^ [BioProject PRJNA486372] using the same parameters for each tool. In the case of cris.py we have pre-merged the reads since it does not work with separated pair-end reads.

### Template-based and substitutions count with CRISPR-A, CRISPR-GA, and CRISPResso2

We have used the previously described simulated data set adding 1100 reads with different template-based modifications to compare the performance of these two tools. We have used a custom python script to generate templates and modified reads with diverse objective modifications including substitutions, insertions, deletions, and delins with different distances to the cut site.

CRISPR-A template-based edits search is based on CIGAR as well as indels search. Although, in this particular case, the user wants to quantify a certain change done with HDR, PE, BE, or any other CRISPR-based technology capable of generating precise edits. For that reason, the quantification is done using the given template and the other counters are updated taking into account the nature of the change. First, the reference is modified using the template that should be shaped by the modification and homology arms in both sites of the change. Then, the new reference with the modification incorporated is aligned against the template with the same algorithm and parameters used to align the sample reads.

Finally, when there are indels in the template, the reads with the same CIGAR are considered as template-based edits. When the template contains substitutions, these are collected taking into account the location and nature of the substitutions.

CRISPR-GA web application was used to quantify the template based simulated data sets using the same templates with different homology arms sizes as used with CRISPR-A, while in the case of CRISPResso2 what we have used is the amplicon reference with the modification integrated with the template. To quantify the template-based reads, as well as base edited reads, with CRISPResso2 we used the same docker as for indels quantification. In the case of prime editing analysis, we get an error that makes us move to the more recent version of CRISPResso2 (2.2.6) that only could be installed with conda in a python3 environment.

### Manual curation of 30 edited samples

We have classified aligned reads as wt or indels with human assessment of 30 different samples from HCT116 edited cell lines [BioProject PRJNA326019]. The 10 targets selected were those with higher edition differences between analyzing using or not a mock for correction. First, sequences with no indels, after alignment of sequencing reads against amplicon reference with minimap2, were classified as wild type. The counts of aligned reads with indels were recalculated using a size bias correction model based on spike-in controls empiric data. Reads with one indel were classified as indels from edition or as wild type, assuming that the indel came from noise, taking into account its length and position respective to the cutting site. Those indels that were not above the cleavage site were manually inspected to decide if their more probable origin was noise or edition. A similar approach was followed for reads with more than one indel. In this case, the main question was if at least one of the indels was probably caused by the editing machinery.

### Exploration of trends in noise

The feature distributions were obtained after comparing the indels detected in 288 control samples, with the ones detected in 864 samples from 3 different cell lines. The comparisons were done by cell line and biological replicate, being this 96 vs 96 samples to not incur on size bias.

### Subtraction performance assessment

All the samples had equal sized amplicons that had their cut site at position 47, allowing stacking and position assessment. To evaluate the performance of the model, we define the parameters of representativity [3] and efficiency [4]. The formal definitions of these parameters are stated below. All these parameters are calculated by position.

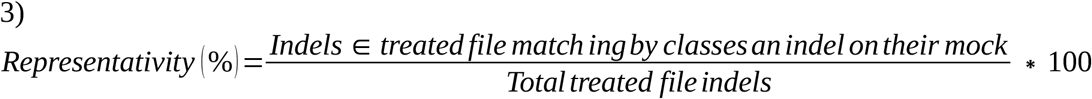

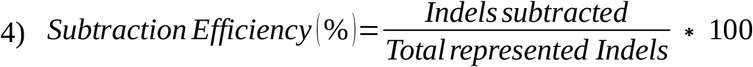

To calculate these, we applied the subtraction algorithm without normalizing, normalizing, and inverse. In other words, comparing the edited file with the normalized mock and not the other way around. To consult the code, read “Data and materials availability”.

### Noise subtraction comparison with ampliCan and CRISPECTOR

ampliCan required the primers used to amplify the amplicon. Since in the data from Van Overbeek et al. paper^26^ [BioProject PRJNA326019] they were not specified, we assumed 15 nucleotides from the start and end of the given amplicon sequences. Some of these sequences were substituted by their reversed complement to allow matching of the reads for ampliCan characterization. Otherwise, the tool does not report edits. Finally, to perform the comparison, the column “Control” from its config file was set to FALSE as it is optional.

As for CRISPECTOR, since it requires a mock file to perform its analysis, fake mock files were generated with the same number of reads reported by this tool for their respective real mock file. In these files, for each sample, all reads correspond to a sequence identical to the provided amplicon so they can be considered mocks free of noise events. From the different resulting parameters, we used the number of edited reads as a reference for both tools.

### HEK293, HCT116, and K563 data analysis

96 targets from 3 different cell lines with three biological replicates per target, with genomic extraction, 48 hours after transfection were analyzed with CRISPR-A analysis pipeline^26^ [BioProject PRJNA326019]. CRISPR-A returns a table with all reads characterized with indels and descriptors of the indels characteristics: kind of modification, position, length, read identifier, micro-homology patterns, nucleotides before the insertion, inserted nucleotides, and nucleotides after the insertion. The content of these tables has been processed with custom R scripts to visualize and explore the results.

RNAseq differential expression analysis of samples from BioProject PRJNA208620 and PRJNA304717 was performed using nf-core/rnaseq pipeline^52^.

### Cloning and plasmids

pDNA-Lama2-gR271 plasmid was performed by Golden Gate assembly using BsaI enzyme and standard methods, by fusing pDNA-BsaI plasmid (vector with Ampicillin resistant gene, the origin of replication derived from the puc19 plasmid and a cloning cassette with BsaI cloning sites) and the Lama2-271-BsaI amplified from C2C12 gDNA.

Different mutations were introduced into pDNA-Lama2-gR271 by site directed mutagenesis following Q5® Hot Start High-Fidelity 2X Master Mix (NEB). Primers were designed to achieve the following indels: 50 nt deletion from 243 nucleotides away from the first amplified nucleotide for targeted sequencing (50_del), deletion of 5 nucleotides from position 277 (5_del), deletion of 1 nucleotide from position 281 (1_del), deletion of 28 nucleotides from position 243 (28_del), deletion of 39 nucleotides from position 243 (39_del), 1 nucleotide insertion at position 281 (1_ins), 5 nucleotides insertion at position 281 (5_ins), and 10 nucleotides insertion at position 281 (10_ins).

### Cell culture, transfection, and electroporation

C2C12 cell line (ATCC CRL-1772) and HEK293T cell lines (Thermo Fisher Scientific) were cultured at 37 °C in a 5% CO2 incubator with Dulbecco’s modified Eagle medium, supplemented with high glucose (Gibco, Thermo Fisher), 10% fetal bovine serum, 2 mM glutamine and 100 U penicillin/ 0.1 mg/ml streptomycin. Cell lines were purchased with an authentication report prior to purchase.

C2C12 electroporation experiment was carried out by using SE Cell Line 4D-Nucleofector (Lonza) and using the manufacturer’s instructions for 100 ul single Nucleocuvette on the 4D-Nucleofector (Lonza). Plasmid DNA molarity ratio was 0.29 gRNA-297-Lama2 : 0.7 Cas9 or GFP.

Lipofectamine 3000 reagent was used to perform the HEK293T transfections following the manufacturer instructions. 240.000 cells were seeded the day before transfection day on a p12 well plate. Plasmid DNA molarity ratio was 0.2 gRNA-TRAC : 0.08 Cas9.

Genomic DNA was extracted using DNeasy Blood and tissue kit (Qiagen) 48 hours post-transfection.

### Library prep and Illumina sequencing for targeted insertion analysis

We implemented Illumina sequencing to capture the modified site with high sensitivity. Plasmids were digested with NotI-HF (NEB) and NheI-HF (NEB) at 37ºC for 1h and inactivated at 80ºC for 20min; the »750 bps was gel purified using the QIAquick Gel Extraction Kit (Qiagen). A qPCR was performed comparing the CTs of 3727 molecules of edited gDNA and the CTs of the same amount of edited DNA supplemented with different amounts of modified plasmids. Two nested PCRs were performed to the mix that had one lower CT number; using KAPA HiFI DNA Polymerase following manufacturer protocol. We did three different cycles on PCR1 (x25, x30 & x35) and two different concentrations in parallel. PCR1 was done with UMIs-Lama2-271-400_fw and UMIs-Lama2-271-400_rv in a 25 ul final volume, and PCR2 with NEB Index primers and NEB Universal Illumina primers. PCR products were purified with QIAquick PCR Purification Kit (Qiagen), mixed in equimolar ratio, and sequenced with Illumina Miseq Nano kit v2.

### Library prep and Illumina sequencing with Unique Molecular Identifiers (UMIs)

Two nested PCR were performed targeting both the 271-Lama2 and TRAC target sites. The Custom PCR UMI (with SQK-LSK109), version CPU_9107_v109_revA_09Oct2020 (Nanopore Protocol) was followed from UMI tagging step to the late PCR and clean-up step. Primers for the UMI tagging were designed using UMI tagging and gene specific primers with amplicon length of around 400 bps. Primers for the early and late PCR were the Illumina Universal primer and Illumina barcoded primer 2 and 4, for 271-Lama2 and TRAC respectively (Table S5). The purified amplicons were then mixed in equimolar ratio and sequenced with Illumina Miseq Nano kit v2.

## Author contributions

Conceptualization: MG, MS-G, AG-V, AS-M

Methodology: MG, AS-M, MS-G, AG-V

Investigation: JJ-W, MS-G, AG-V, SJ, ME, EB

Software: MS-G, AG-V, SJ, ME, EB

Visualization: MS-G, AG-V, SJ

Formal analysis: MS-G, AG-V, SJ

Supervision: MG, AS-M

Writing—original draft: MS-G, AG-V, SJ, JJ-W

Writing—review & editing: MS-G, AG-V, SJ, EB, MG

## Data and materials availability

Next-generation sequencing data are available in the European Nucleotide Archive under the Study accession number PRJEB53901. SimGE developed R package can be installed with devtools: devtools::install_bitbucket(“synbiolab/SimGE”). Code for CRISPR-A pipeline has been made available in Bitbucket https://bitbucket.org/synbiolab/crispr-a_nextflow/ and through the web page application https://synbio.upf.edu/crispr-a/. This pipeline will also be added to the NF-core community. Custom analysis scripts for data analysis and visualization are freely available at https://bitbucket.org/synbiolab/crispr-a_figures/.

